# Deciphering histone mark-specific fine-scale chromatin organization at high resolution with Micro-C-ChIP

**DOI:** 10.1101/2023.10.30.563558

**Authors:** Mariia Metelova, Nils Krietenstein

**Affiliations:** Novo Nordisk Foundation Center for Protein Research, University of Copenhagen

## Abstract

The regulation of cell-type-specific transcription programs is a highly controlled and complex process that needs to be fully understood. The gene regulation is often influenced by distal regulatory elements and their interactions with promoters in three-dimensional space. Although proximity ligation techniques like Hi-C have revolutionized our understanding of genome organization, the genomic resolution for many of these methods is limited by both experimental and financial constraints. Here, we introduce Micro-C-ChIP to provide extremely high-resolution views of chromosome architecture at genomic loci marked by specific covalent histone modifications. This is achieved by chromatin immunoprecipitation of specific chromatin states to target chromosome folding libraries to focus on chromatin domains (regulatory elements, heterochromatin, etc.) of interest, yielding extremely high sequencing depth at these loci. We applied Micro-C-ChIP to mouse embryonic stem cells (mESC) and hTERT-immortalized human retinal epithelial cells (hTERT-RPE1), revealing architectural features of genome organization with comparable or higher resolution than Micro-C datasets sequenced with higher depth. We discovered extensive promoter-promoter networks in both cell types and characterized the specific architecture of bivalently marked promoters in mESC. Together, these data highlight Micro-C-ChIP as a cost-effective approach to exploring the landscape of genome folding at extraordinarily high resolution.

## Introduction

The topological organization of genomes within the nucleus is consequential for many DNA-templated processes such as gene transcription^1–5^, DNA damage repair^6,7^, and DNA replication^8^. The development of sequencing-based approaches to 3D genome architecture, such as Hi-C, has revolutionized how we view genome organization^9–11^. Hi-C is based on Chromosome Conformation Capture (3C), in which chromatin is crosslinked, digested with restriction enzymes (RE), and ligated to generate chimeric DNA molecules from topologically proximal regions. At the scale of chromosomes, Hi-C has revealed that active and inactive eu- and heterochromatin are spatially separated into A and B compartments^9^. At the level of genes, topologically associating domains (TADs) are formed by loop-extrusion machinery to regulate the interaction probability of genomic elements, such as promoters and enhancers, in 3D space^11–16^. Enhancers are distal regulatory elements that modulate the transcriptional activity of promoters, presumably via direct spatial interaction^17^.

However, with the resolution of Hi-C, detecting focal interactions remains challenging^18^. For instance, resolution in Hi-C is limited by the choice of restriction enzymes and the heterogeneous spacing of their target sites and sequencing required for maximal resolution (>1 billion reads) can be extremely expensive. Micro-C, an MNase-based version of Hi-C, improves the detection of short-range features such as enhancer-promoter loops by utilizing a dual crosslinking strategy and nucleosome resolution fragmentation^19–22^. Moreover, the recent development of locus-specific capture strategies allows much deeper sequencing of a locus of interest, highlighting the importance of resolution for understanding how genomes are folded^23^. These approaches have highlighted novel features like micro-compartments and resolved the dynamic changes in topological interactions in response to the depletion of important transcriptional factors. However, despite their extremely high resolution, such sequence-based capture strategies are limited to a small region in the genome.

Chromatin immunoprecipitation (ChIP)-enrichment strategies of Hi-C-like protocols, such as HiChIP^24^, PLAC-seq^25^, and ChIA-PET^25,26^, have proven to be valuable tools to overcome the extremely high burden of sequencing costs. Here, we combine the high-resolution method Micro-C with histone PTM-specific chromatin immunoprecipitation, a hybrid methodology we call Micro-C-ChIP. Because Micro-C leverages MNase as the chromatin fragmenting enzyme, which digests accessible DNA and leaves nucleosomes intact, this strategy is ideal for determining the 3D interactions of genomic regions specifically marked by histone post-translational modifications (PTMs). Here, we resolved the 3D genome organization specifically at either H3K4me3- or H3K27me3-marked domains, using Micro-C-ChIP in mESC and hTERT-RPE1 cells. While H3K4me3 is associated with transcription start sites (TSS)^27,28^ and has been recently shown to be involved in regulating RNAPII pause-release in mESC^29^, H3K27me3 is a repressive histone modification^30,31^. In pluripotent cells, bivalent promoters are simultaneously marked by both repressive H3K27me3 and active H3K4me3 marks^32,33^ in contrast to differentiated cells where both marks are largely exclusive^34^. We uncover an extensive promoter-originating connectome in differentiated and pluripotent cells at active promoters. Surprisingly, bivalently marked promoters adopt structures of active genes at transcription start sites, implicating the 3D genome organization in the readiness to rapidly respond to environmental queues during differentiation to become transcriptionally active or repressed.

### Micro-C-ChIP development: a tool to detect histone mark-specific 3D genome interactions

To explore chromosome folding across chromatin domains marked with specific PTMs, we set out to carry out Micro-C to map chromatin organization of genomic loci isolated using PTM-specific antibodies. We modified and optimized the existing Micro-C protocol, carrying out immunoprecipitation of specific histone marks after the proximity ligation step (**Fig. 1a**). Briefly, nuclei derived from dually crosslinked mouse embryonic stem cell (mESC) were MNase-digested, the DNA ends were biotin-labeled, and proximity ligated. Ligated chromatin was sonicated to solubilize the heavily crosslinked chromatin. The optimal conditions, e.g., sonicator, sonication cycles, and concentration of detergent, were selected to release a high fraction of proximity-ligated di-nucleosomal-sized DNA fragments into the soluble fraction (**Fig. 1b**). For this proof-of-concept study, we immunoprecipitated two histone marks, H3K27me3 and H3K4me3.

**Fig.1:**
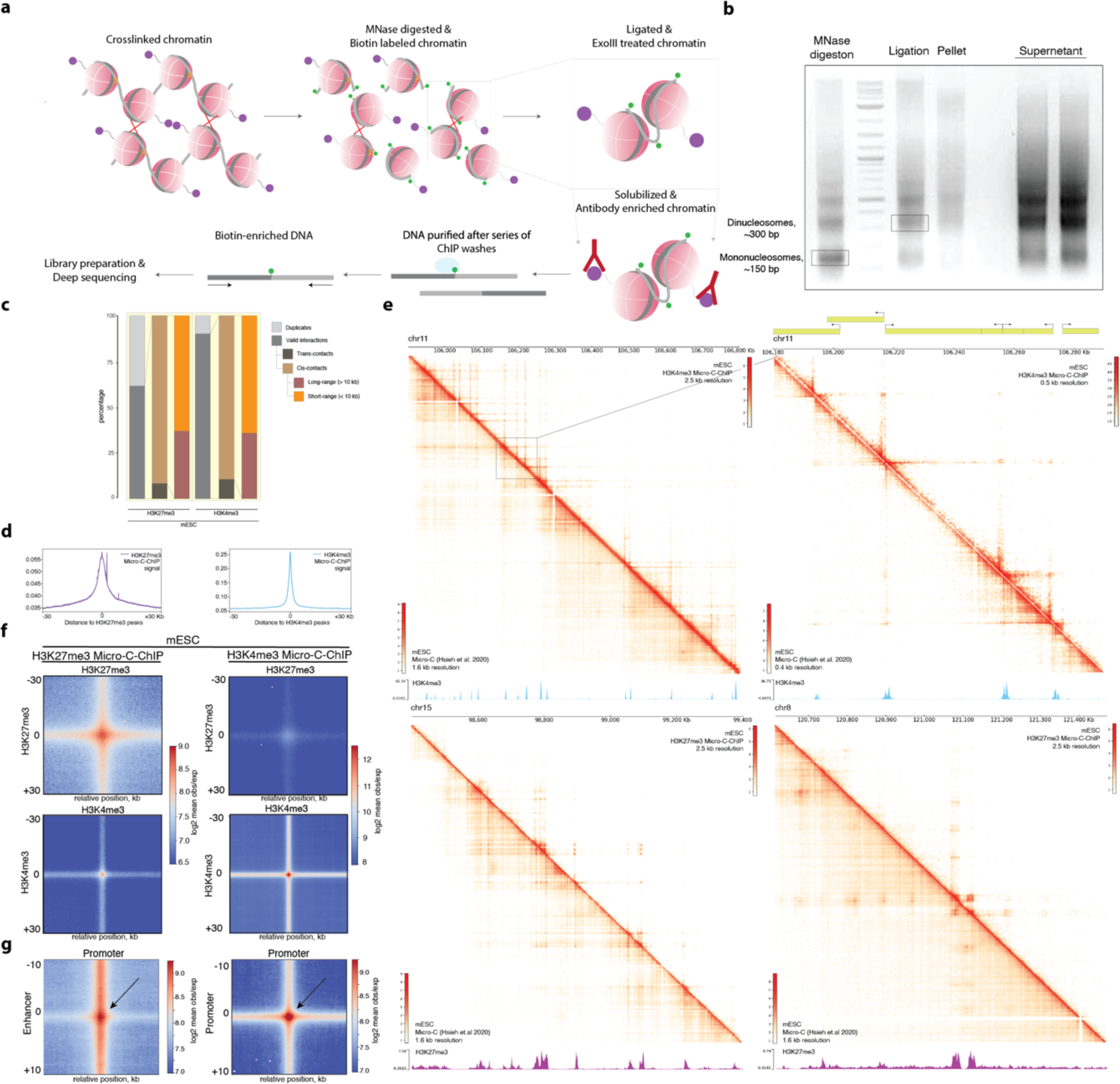
Micro-C-ChIP detects histone mark-specific 3D genome interactions in mESC. **a**, Outline of the Micro-C-ChIP method. The cells are dually crosslinked, derived nuclei are MNase-digested to mono-nucleosomal fragments which are then biotinylated, ligated and ExoIII treated. The sonication is performed to solubilize heavily dually crosslinked chromatin and to perform immunoprecipitation against histone mark of interest. Antibody-bound ligated fragments are then washed, DNA is purified and pulled-down with streptavidin and sequencing library is prepared. **b**, 1.5% Agarose gel electrophoresis of digested chromatin (5U MNase, Lane1), compared to proximity-ligated sample (Lane 3), and sample pellet (Lane 4) and supernatant (Lane 5) after sonication. Lane 2: NEB 2-log DNA ladder. **c**, Sequencing statistics of the merged mESC H3K27me3 and H3K4me3 Micro-C-ChIP datasets. **d**, CPM-normalized signal of H3K27me3 and H3K4me3 Micro-C-ChIP datasets at the respective ChIP-seq peak sites. **e**, Unbalanced interaction heatmaps comparing Micro-C-ChIP (top-right triangle) to bulk Micro-C (bottom-left triangle) (Hsieh et al., 2020). Top panels: H3K4me3 interaction maps at 2.5 kb and 0.5 kb resolution. Bottom panels: H3K27me3 interaction maps at 2.5 kb resolution. Genome coordinates and gene annotations are shown above and corresponding ChIP-seq tracks (Supplementary Table 1) are shown below respective panels. **f**, Off-diagonal pile-up of H3K27me3 and H3K4me3-enriched Micro-C-ChIP datasets (500 bp resolution) at loci of paired ChIP-seq H3K27me3 (upper panel) and H3K4me3 (lower panel) peaks. The intersected ChIP-seq peaks at the distance between 10 and 300 kb were used. **g**, Pileup plots of H3K4me3 Micro-C-ChIP (200 bp resolution) at putative enhancer-promoter (E-P) (left) and promoter-promoter (P-P) (right) interactions at ±10kb window.

**Extended Data Fig. 1.**
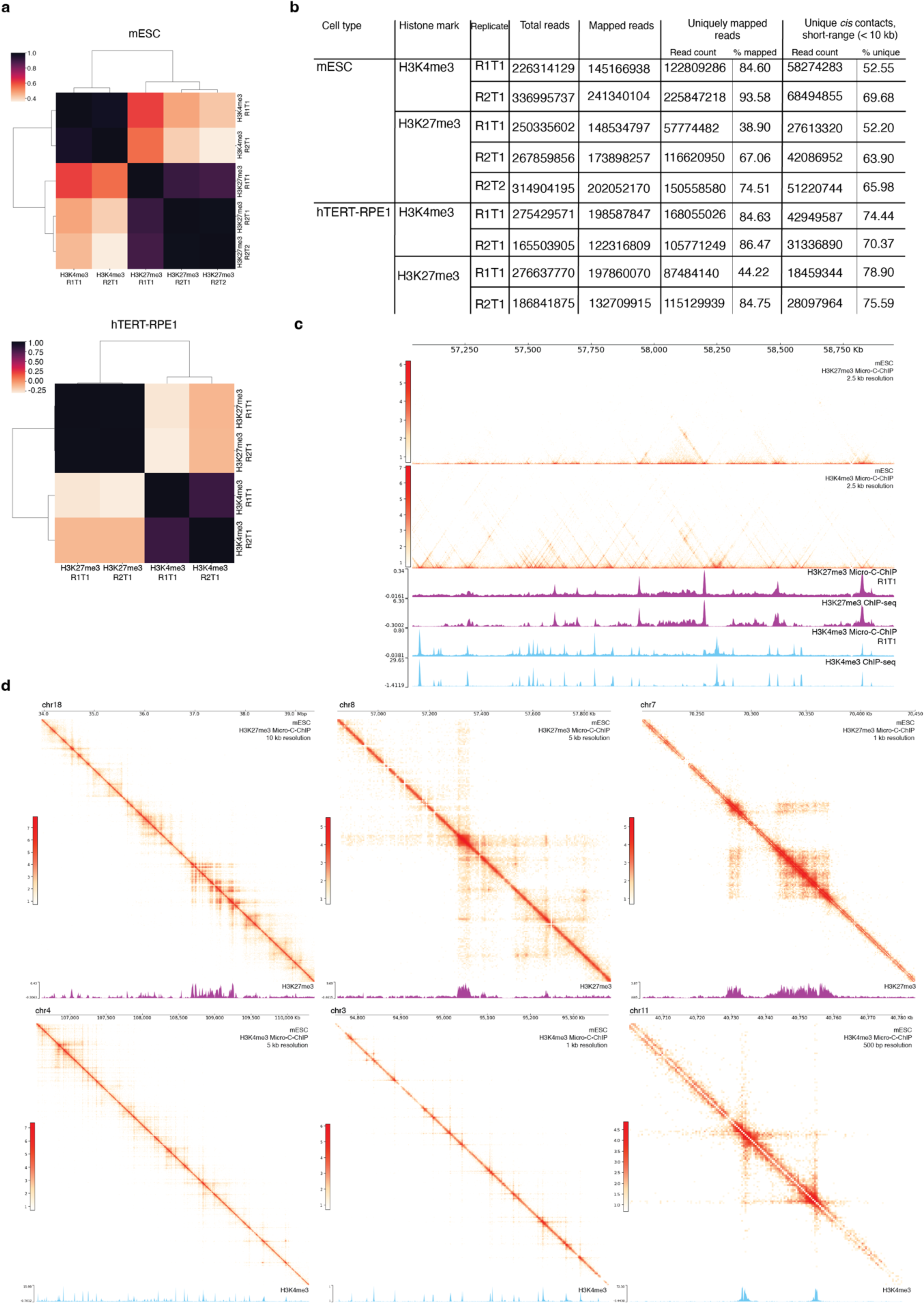
Micro-C-ChIP reproducibly captures highly precise histone mark-specific interactions. **a**, Heatmaps of reproducibility scores for all replicates across both H3K4me3- and H3K27me3-enriched Micro-C-ChIP datasets in mESC (top) and hTERT-RPE1 (bottom). Reproducibility scores were determined using HiCRep at 10 kb resolution. **b**, Sequencing statistics of all Micro-C-ChIP replicates. The numbers of the total and mapped sequencing reads, unique reads and short-range (< 10 kb) unique *cis*-reads are shown. R1 and R2 indicate biological replicates, T1 and T2 - technical replicates. **c, d,** Unbalanced interaction heatmaps of merged H3K27me3 (top) and H3K4me3 (bottom) Micro-C-ChIP mESC datasets at different resolutions. CPM-normalized Micro-C-ChIP signal tracks for single replicates (c) and ChIP-seq signal tracks for the corresponding histone mark (c and d) are shown below the contact maps.

Biological and technical replicates for H3K4me3 and H3K27me3 Micro-C-ChIP were visually indistinguishable, with correlation coefficients ranging from ∼0.8 to 0.9x (**Extended Data Fig. 1a**). Therefore, replicate datasets were merged for downstream analysis. In total, the combined replicates yielded approximately 300 million valid read pairs for each condition (**Extended Data Fig. 1b**). A high ratio of non-duplicated reads for H3K27me3 and H3K4me3 (62% and 90% respectively) confirmed the quality of Micro-C-ChIP libraries (**Fig. 1c**). Among valid pairs, intra-chromosomal (cis) interactions comprised 92% of all contacts for H3K27me3 Micro-C-ChIP, and 89% for H3K4me3 Micro-C-ChIP. *Cis*-interactions between loci separated by less than 10 kb prevailed in H3K27me3 and H3K4me3 Micro-C-ChIP datasets, comparable to the bulk Micro-C yields^18^.

To confirm the enrichment in individual replicates, we visualized the Micro-C-ChIP 1D signal tracks for both histone marks compared to published ChIP-seq datasets (**Extended Data Fig. 1c**). The Micro-C-ChIP coverage tracks align with ChIP-seq profiles by visual inspection, which is also corroborated by plotting the average signal of Micro-C-ChIP datasets at the corresponding ChIP-seq peaks (**Fig. 1d**). To evaluate the performance of Micro-C-ChIP further, we visually compared contact heatmaps for Micro-C-ChIP against H3K27me3 and H3K4me3 marks with the deeply sequenced Micro-C mESC dataset from Hsieh et al^1^. The Micro-C-ChIP matrices for both marks show strong interaction signals at regions marked by the respective histone mark from published ChIP-seq data (**Fig. 1e, Extended Data Fig. 1c, d**). H3K4me3 ChIP-peaks are characterized by narrow, precise peaks at the promoter regions compared to wider H3K27me3 ChIP-peaks, and these patterns are translated onto the 3D genome landscape. Here, H3K4me3 Micro-C-ChIP reveals fine-tuned precise contacts forming a grid-like structure, whereas H3K27me3 Micro-C-ChIP displays 3D-structural patterns with both stripe- and block-like structures. Importantly, despite the much lower sequencing depth, Micro-C-ChIP detects structural features of bulk Micro-C with higher definition.

Promoter-promoter interactions can be visualized by H3K4me3-to-H3K4me3 intersections^1^. These intersections show a focal enrichment at the center of the interaction pile-up plot and stripes emerging from the promoter sites, generating a cross-pattern. These patterns are generated by active loop-extrusion (LE)^12,13^. Here, loop-extrusion factors, such as cohesin, extrudes chromatin bi-directionally until it is stalled. When the LE is stalled on one side, it extrudes chromatin uni-directionally. Consequently, the stalled site is brought into close contact with many other genomic regions visible as stripes in interaction heatmaps and pile-up plots. Similarly, two genomics loci are kept spatially close if LE is stalled completely. To validate the enrichment of Micro-C-ChIP libraries at expected chromatin domains, we computed the intersections of H3K27me3 with H3K27me3 and H3K4me3 with H3K4me3 from available ChIP-seq peaks. We plotted the Micro-C-ChIP signal for both histone PTMs at both intersection sites (**Fig. 1f**), each showing strong focal enrichment at the center of the plot and emerging stripes (H3K27me3: top-left panel, H3K4me3: bottom-right panel), which confirms the enrichment of histone mark-specific interactions. In addition, we observe weaker but similar signals at non-corresponding interaction sites, e.g., H3K27me3-Micro-C-ChIP plotted at H3K4me3 intersections (**Fig. 1f**, bottom-left panel) and vice-versa (top-right panel). We suspect that the overlapping signal reflects bivalent chromatin, marked by the active H3K4me3 and repressive H3K27me3 histone PTMs.

To test whether Micro-C-ChIP preferentially enriches sites that are marked by histone PTMs at both anchors, we plotted the observed/expected signal at putative enhancers with promoters (E-P) and promoters with promoters (P-P) intersection sites (**Fig. 1g**). In bulk Micro-C experiments, the focal signals at these sites are comparable^1,4^. In H3K4me3 Micro-C-ChIP, we also observe similar focal intensities at P-P and E-P intersection sites. This confirms that P-P interactions, which are enriched for H3K4me3 at both anchors, are not preferentially enriched over E-P interactions, which are only marked by H3K4me3 at the promoter site. We conclude that Micro-C-ChIP yields histone modification-specific proximity-ligation libraries with a high signal-to-noise ratio at moderate sequencing depth.

### Micro-C-ChIP reproducibly works in a different cell type

To test if the overlapping signal of H3K4me3 and H3K27me3 Micro-C-ChIP reflects bivalently marked promoters, we generated Micro-C-ChIP libraries for the same histone marks from differentiated hTERT-immortalized human RPE1 cells (hTERT-RPE1). Here, the libraries were sequenced to at least 200 million mapped valid read pairs (**Extended Data Fig. 1a, b**), and the resulting dataset was processed as described for mESC. Similar to mESC, H3K27me3 yielded a lower non-duplicated rate (61%) compared to H3K4me3 (85%) (**Fig. 2a, Extended Data Fig. 1a, b**). Again, the distribution of long vs. short-range reads are comparable to published bulk Micro-C datasets, confirming the overall quality and reproducibility of the Micro-C-ChIP protocol.

**Fig.2:**
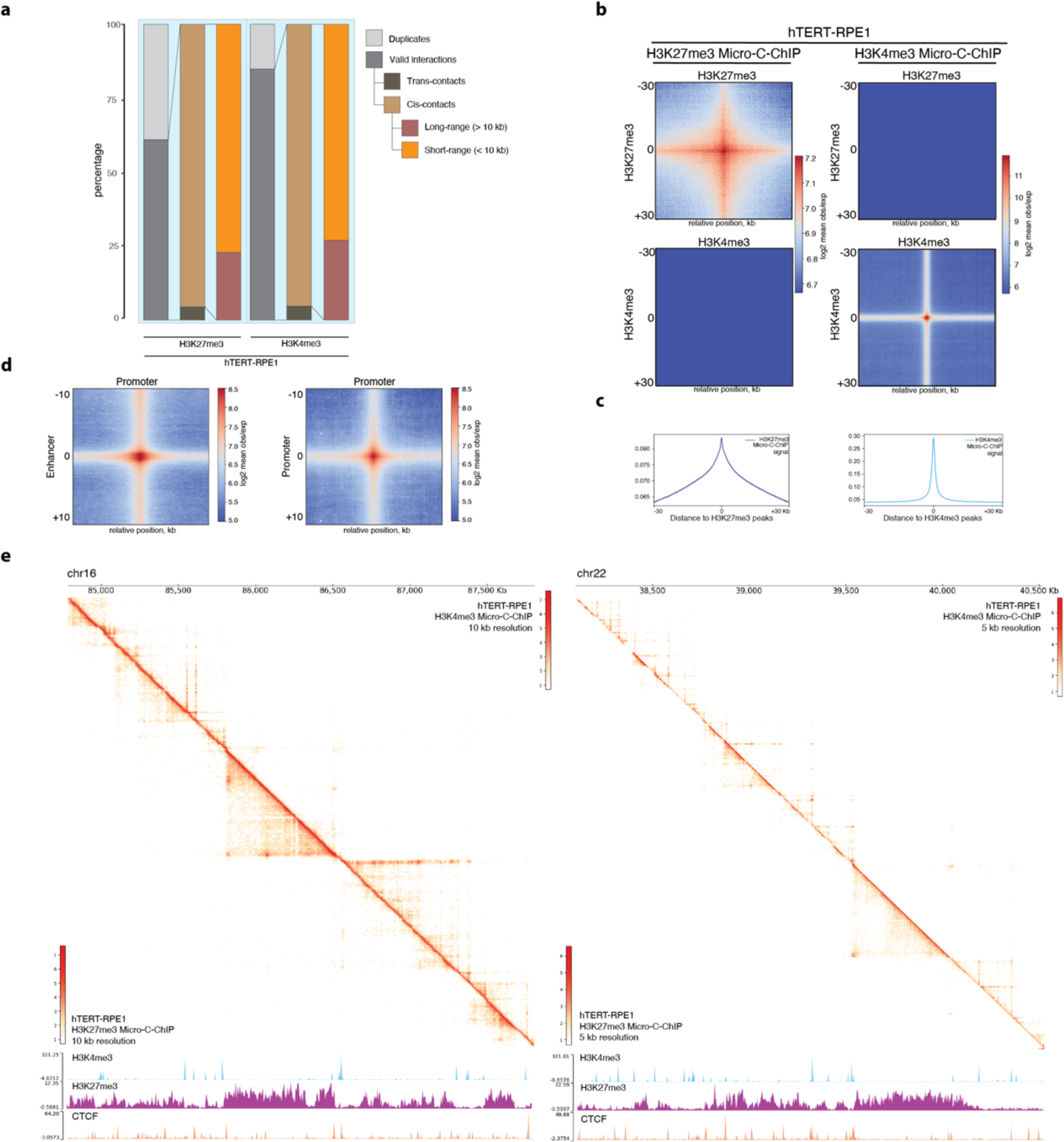
Micro-C-ChIP is applicable in differentiated hTERT-RPE1 cells. **a**, Sequencing statistics of merged hTERT-RPE1 H3K27me3 and H3K4me3 Micro-C-ChIP datasets. **b**, As Fig. 1e but for hTERT-RPE1 cells. **c**, CPM-normalized 1D signal computed from H3K27me3 and H3K4me3 Micro-C-ChIP datasets at the ChIP-seq peaks. **d**, Pileup plots of H3K4me3 Micro-C-ChIP (200 bp resolution) at putative enhancer-promoter (E-P) (left) and promoter-promoter (P-P) (right) interactions within ±10kb **e**, Unbalanced interaction heatmaps comparing H3K27me3 (bottom-left triangle) and H3K4me3 (top-right triangle) Micro-C-ChIP at 5 kb (right panel) and 10 kb (left panel) resolution. Genome coordinates are show above and ChIP-seq signal tracks for H3K4me3, H3K27me3, and CTCF are shown below the contact maps.

**Extended Data Fig. 2.**
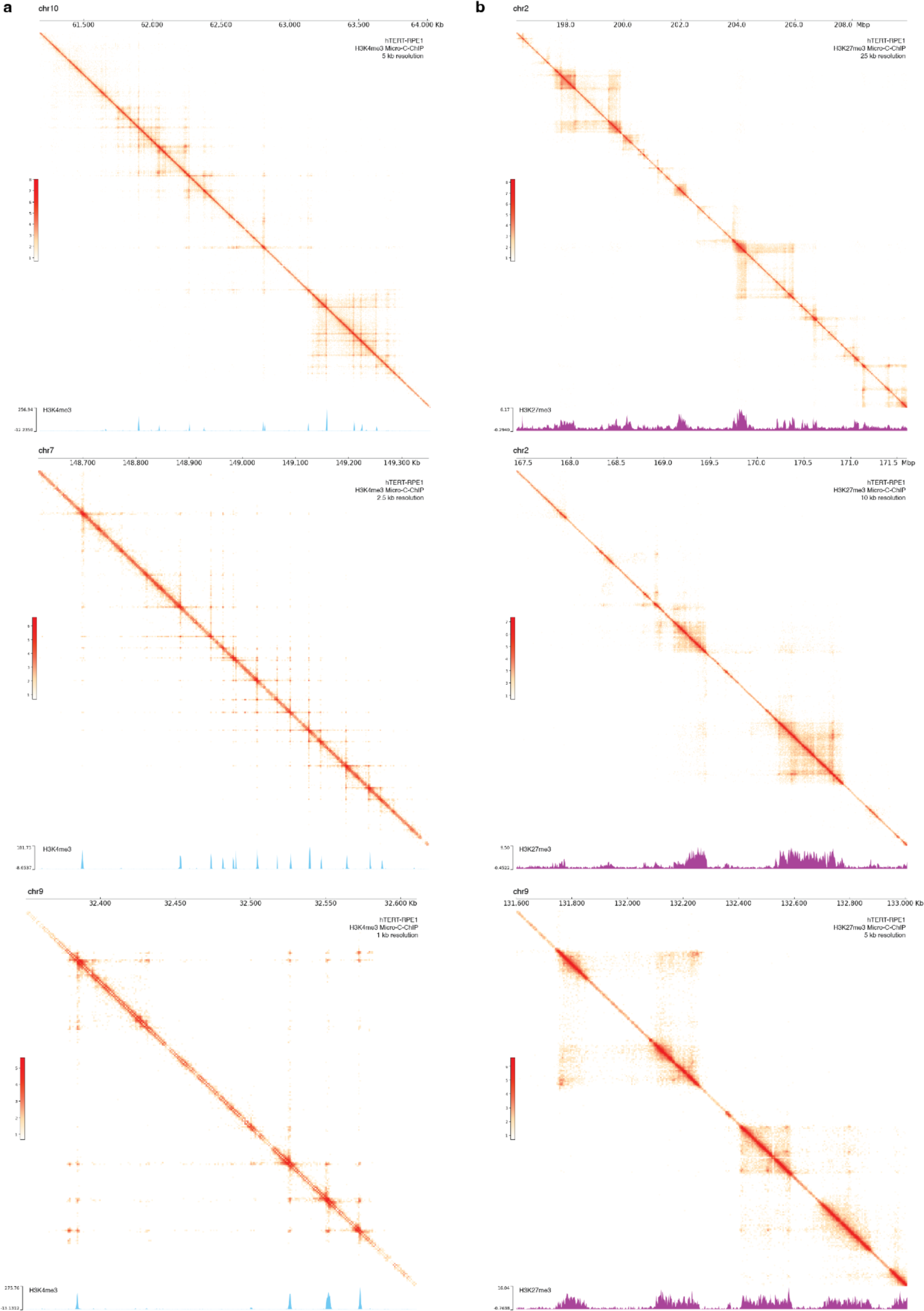
Detection of histone modification-specific interactions in hTERT-RPE1 cells. **a**, **b,** H3K4me3-(a) and H3K27me3 (b)-enriched contact maps of hTERT-RPE1 cells at different resolutions.

To confirm the absence of bivalent chromatin regions, we plotted the interactions measured with Micro-C-ChIP at the intersections of H3K27me3 peaks or H3K4me3 (**Fig. 2b**). Again, H3K27me3 displays strong interactions at H3K27me3 ChIP-seq peaks but, in contrast the mESC (**Fig. 1f**), no signal at H3K4me3 intersected ChIP-seq peaks. Similarly, H3K4me3 interactions are enriched at H3K4me3 but not at H3K27me3-enriched intersected loci. The successful enrichment was also confirmed by plotting the average CPM-normalized signal of Micro-C-ChIP datasets at the regions of the corresponding histone mark peaks (**Fig. 2c**). Like mESC (**Fig. 1g**), H3K4me3 Micro-C-ChIP shows similar focal interaction signals putative at P-P and E-P intersection sites in hTERT-RPE1 (**Fig. 2d**).

In contrast to mESC, H3K27me3 and H3K4me3 chromatin marks are more binary in differentiated cells. We visualized Micro-C-ChIP contact matrices across relevant PTM-marked regions. Focusing on genomic loci carrying both histone marks, we confirm that Micro-C-ChIP libraries yield highly distinct patterns for each mark with almost no overlap (**Fig. 2e, Extended Data Fig. 2a, b**). H3K27me3 marks broad regions of inactive chromatin reflected in 3D interaction heatmaps. In contrast to mESC, H3K27me3 displays little to no evidence for interactions “stipes” or “flares”, reflecting the loss of punctate K27me3 and preservation of broad K27 domains that occur as pluripotent cells differentiate. Conversely, H3K4me3 Micro-C-ChIP is characterized by a precise grid-like structure consisting of stripes and dots stemming from the punctate localization pattern of this modification at promoters. From these observations, we conclude that Micro-C-ChIP reproducibly captures 3D organization specifically at domains of interest in different cell types.

### H3K4me3 Micro-C-ChIP identifies widespread promoter-associated loops

To get a better understanding of the functional role of histone mark-specific interactions, we called sites of enriched interactions, apparent as “dots” in contact maps and inferred to represent chromosome loops, for our H3K4me3 Micro-C-ChIP datasets with *Peakachu* pipeline. We detected 72574 loops in mESC and 63085 loops in hTERT-RPE1. Visual inspection showed that most of the called loops align with H3K4me3 ChIP-seq peaks and RNAPII signal in both cell types, suggesting that loops are associated with ongoing transcription (**Fig. 3a**). The computational detection of loops at shorter distances is complicated by increased noise from unspecific interactions close to the diagonal. Micro-C detects chromosome loops at shorter distances more efficiently than Hi-C, indicating an improved signal-to-noise-ratio^18^. To estimate if Micro-C-ChIP improves the detection of these short-range interactions, we plotted the distances of called loops as CDF (**Fig. 3b**). Distances between anchors of H3K4me3 Micro-C-ChIP detect loops are smaller compared to the deepest sequenced mESC dataset available^1^, indicating a higher signal-to-noise, which allows the detection of short-range features, such as loops. Notably, the detection of short-range chromosome interaction is even higher in differentiated hTERT-RPE1.

**Fig.3:**
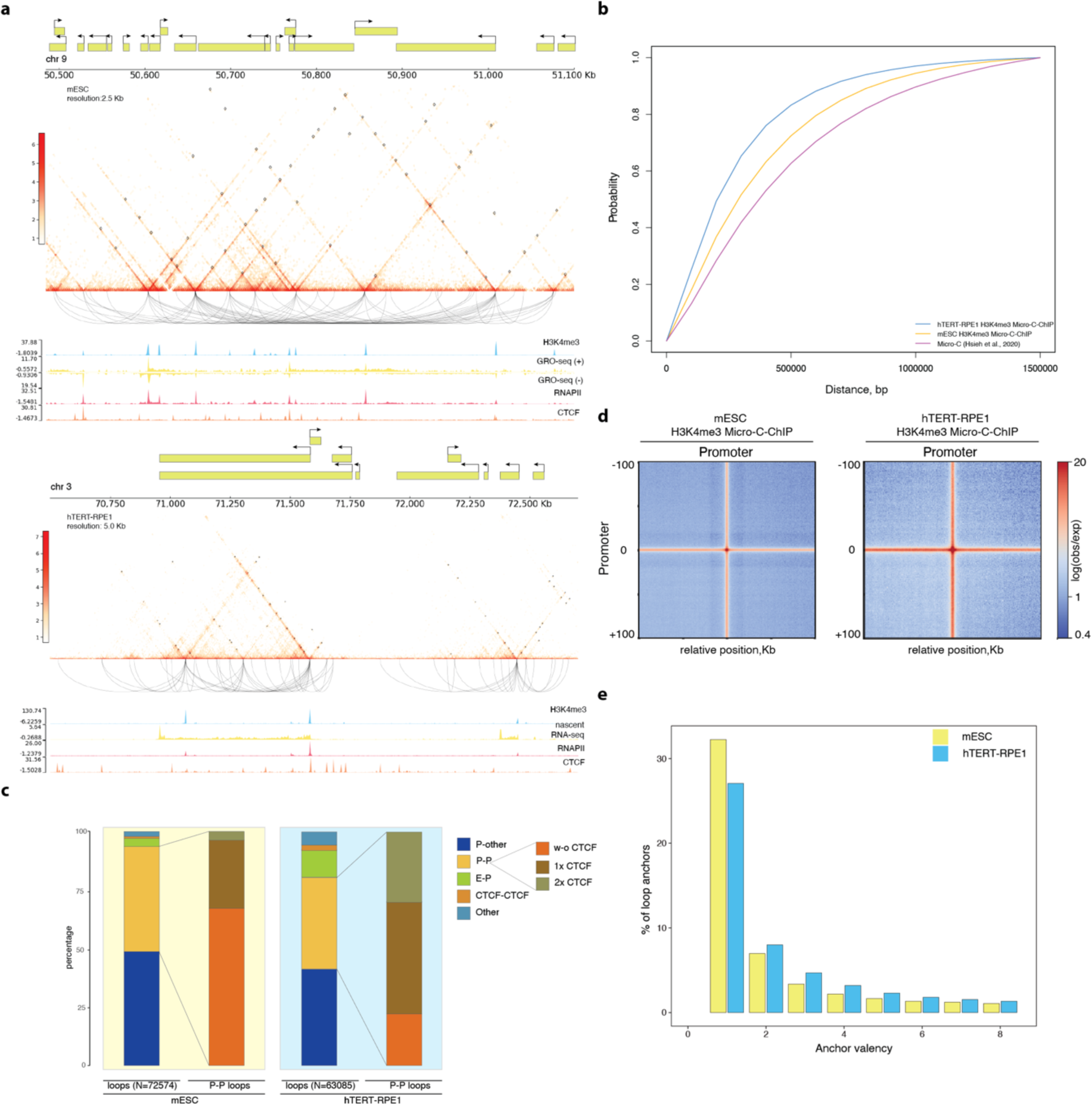
Promoter-associated loops prevail in H3K4me3-enriched Micro-C-ChIP datasets. **a**, Unbalanced interaction heatmaps for mESC (top) and hTERT-RPE1 (bottom) H3K4me3 Micro-C-ChIP at 2.5 kb and 5 kb resolution, respectively. Loops called at 2.5 kb resolution via the Peakachu pipeline are shown on the heatmaps as squares. Loop arcs show enriched interactions. 1D chromatin tracks are shown below the contact maps (Supplementary Table 1). **b**, Cumulative distribution function (CDF) plot of distances between loop anchors in H3K4me3 Micro-C-ChIP in mESC and hTERT-RPE1 and bulk Micro-C (Hsieh et al., 2020). **c**, Percentage of H3K4me3-enriched loops classified into the following categories: P-P (promoter-promoter), P-Other, E-P (enhancer-promoter), CTCF-CTCF and Other. CTCF-CTCF loops include only anchors that do not overlap with any regulatory elements. P-P loops are further classified by presence of CTCF: At one of the anchors (1x CTCF) or both anchors (2x CTCF). The loop categorization is shown for both mESC (left) and hTERT-RPE1 (right). **d**, Interaction pileups at P-P loops (classified in Fig. 3b) for mESC (left) and hTERT-RPE1 (right). Data is shown for ±100 kb window (100 kb upstream and 100 kb downstream) surrounding loop anchors. **e**, Histogram of loops anchor valencies of for mESC and hTERT-RPE1 H3K4me3 Micro-C-ChIP.

**Extended Data Fig. 3.**
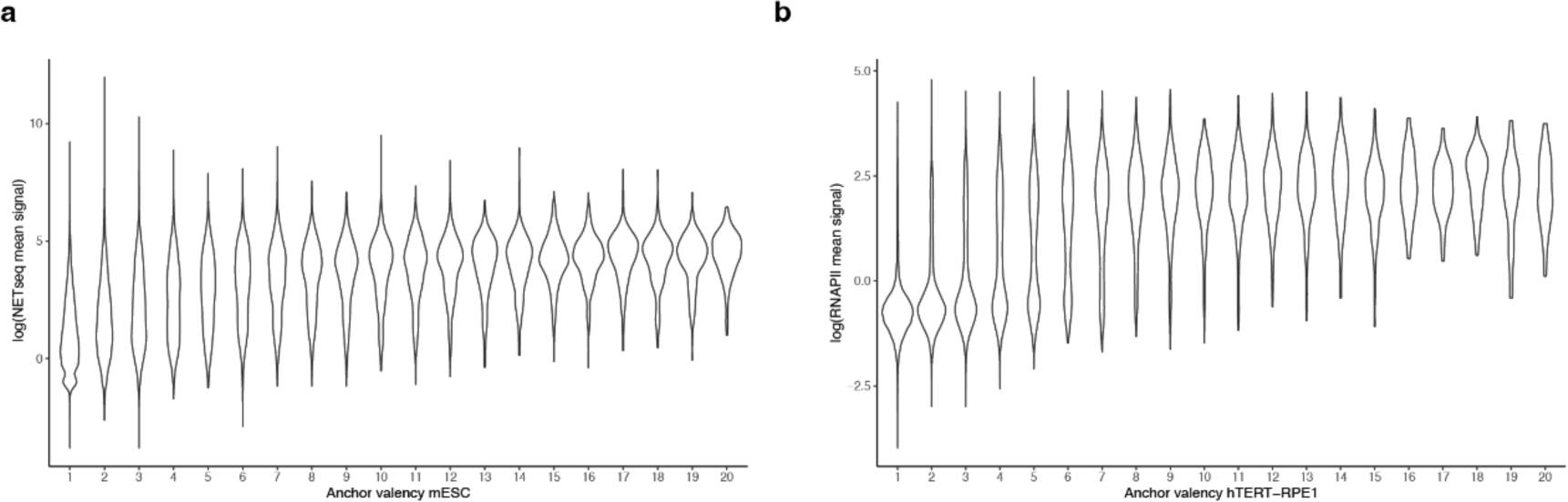
H3K4me3-enriched anchors are engaged in more loops associated with higher transcription rate in both mESC and hTERT-RPE1 cells. **a, b,** Violin plots of log-transformed mean NET-seq (a) and RNAPII (b) signal at loops anchors clustered based on their valency in mESC and hTERT-RPE1, respectively.

To understand which genomic elements are connected by chromosome loops, we classified loop anchors for both cell types into promoter-promoter (P-P), enhancer-promoter (E-P), CTCF-CTCF, promoter-other (P-other), and other interaction sites (**Fig. 3c**). 90-95% of all identified interaction sites were classified as promoter-associated interactions (P-P, E-P, and P-other), which is expected for the promoter mark enriched H3K4me3 Micro-C-ChIP libraries. In mESC, 45% of all loops represent P-P interactions, while only 5 % are classified as E-P interactions. In hTERT-RPE1 cells, a larger fraction of detected interactions connects promoters with enhancers (12%), although most interactions remain between promoters (39%). Notably, manually annotated chromosome loops in RCMC experiments showed an enrichment of P-P over E-P interactions, demonstrating that the detection of P-P interactions is a matter of resolution^23^.

H3K4me3 interactions show intensive networks between promoters. We validated the presence of strong interactions between promoters in mESC and hTERT-RPE1 by plotting averaged contact signals of identified P-P loops for each cell type, (**Fig. 3d**). Interestingly, 60% of all P-P loops were not associated with detectable CTCF in mESC. In contrast, in hTERT-RPE1 cells we find that 78% of all P-P interactions were associated with at least one CTCF peak at either anchor site (**Fig 3c**). Next, we computed the anchor valency, a measurement of how often a given anchor is engaged in chromosome loops (**Fig. 3e**). Despite fewer loops being called in hTERT-RPE1, the valency increases in this differentiated cell type. Together, in comparison to Micro-C, we showed that even at lower sequencing depth (300 million reads vs 3.18 billion filtered reads^1^), Micro-C-ChIP can detect a considerable number of loops with a substantial portion of close-to-diagonal contacts. Moreover, those loops possess distinctive features depending on the cell type. Most importantly, H3K4me3-enriched loops are P-P contacts and anchor valency of promoters largely increases with transcription rate in both cell types (**Extended Data Fig. 3a, b**).

### Transcription is not a prerequisite of stripe formation in mESC

The formation of chromosome loops and stripes are characteristic features of the active process loop extrusion process^35^. Computational modeling proposes that RNAPII functions as a moving barrier for loop-extrusion^5^. Experimental data supports this model and shows that stripes and chromosome loops scale with transcriptional activity^3,4^. Visual inspections of H3K27me3 and H3K4me3 interaction heatmaps show stripes for both marks in mESC, and only for H3K4me3 in hTERT-RPE1 (**Fig. 1d, 2e, 3a, Extended Data Fig. 1c, d, 2a, b**). Putative P-P contacts were recently shown to be unaffected by RNAPII depletion^4^, which contrasts with previous studies^1^. Thus, the effect of transcription on promoter connectome is unclear.

Leveraging the ability to annotate P-P interactions from 3D data, we investigated how P-P contacts identified from the H3K4me3 dataset (**Fig. 3d**) scale with transcription, stratifying them by RNAPII level (**Fig. 4a**). We find that focal P-P contact intensities are comparable between promoters with different RNAPII level, although the stripe intensities scale down with RNAPII level reduction. To test if these P-P contacts also include bivalent promoters, we plotted H3K27me3 Micro-C-ChIP data at these sites. We observe an overlap between H3K27me3 and H3K4me3 signal at RNAPII depleted sites in mESC (**Fig. 4a**). This effect is more prominent in mESC than in hTERT-RPE1 cells. This supports the finding that bivalent promoters, such as the *Hox* gene cluster, engage in larger P-P networks in mESC^36^.

**Fig. 4:**
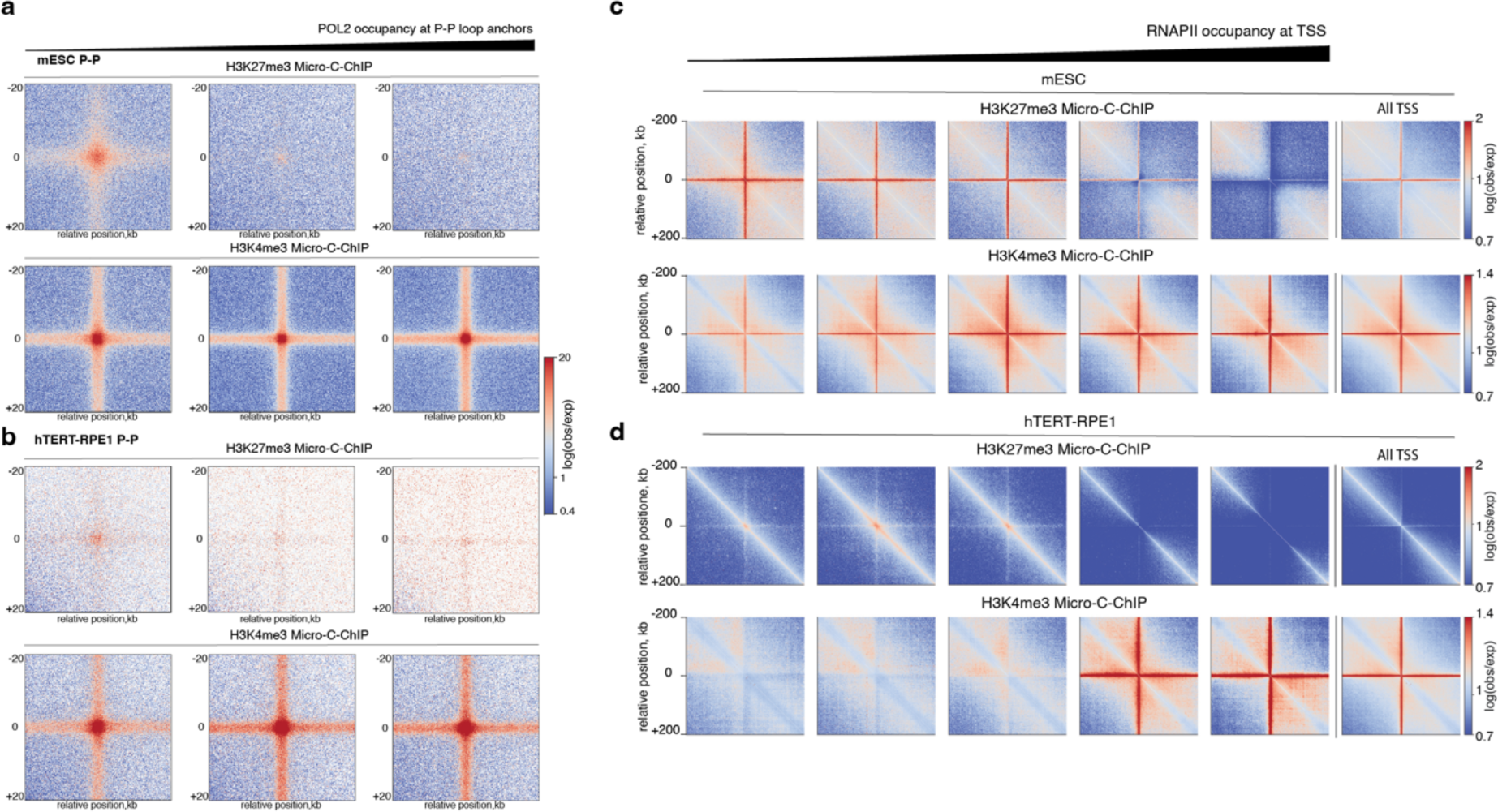
H3K27me3 Micro-C-ChIP reveals 3D signatures of active promoters at repressed TSS exclusively in undifferentiated cells. **a**, H3K27me3 (top) and H3K4me3 (bottom) Micro-C-ChIP interaction pileups at P-P loops from Fig. 3e stratified by RNAPII occupancy at loop anchors into three quantiles for ±20 kb for mESC. **b**, Same as in a, but for hTERT-RPE1. **c**, TSS-centered pileups of H3K27me3 (top) and H3K4me3 (bottom) Micro-C-ChIP in mESC stratified by RNAPII occupancy into five quantiles. The sixth panel represents cumulative interactions of TSS. **d**, Same as c, but for hTERT-RPE1.

**Extended Data Fig. 4.**
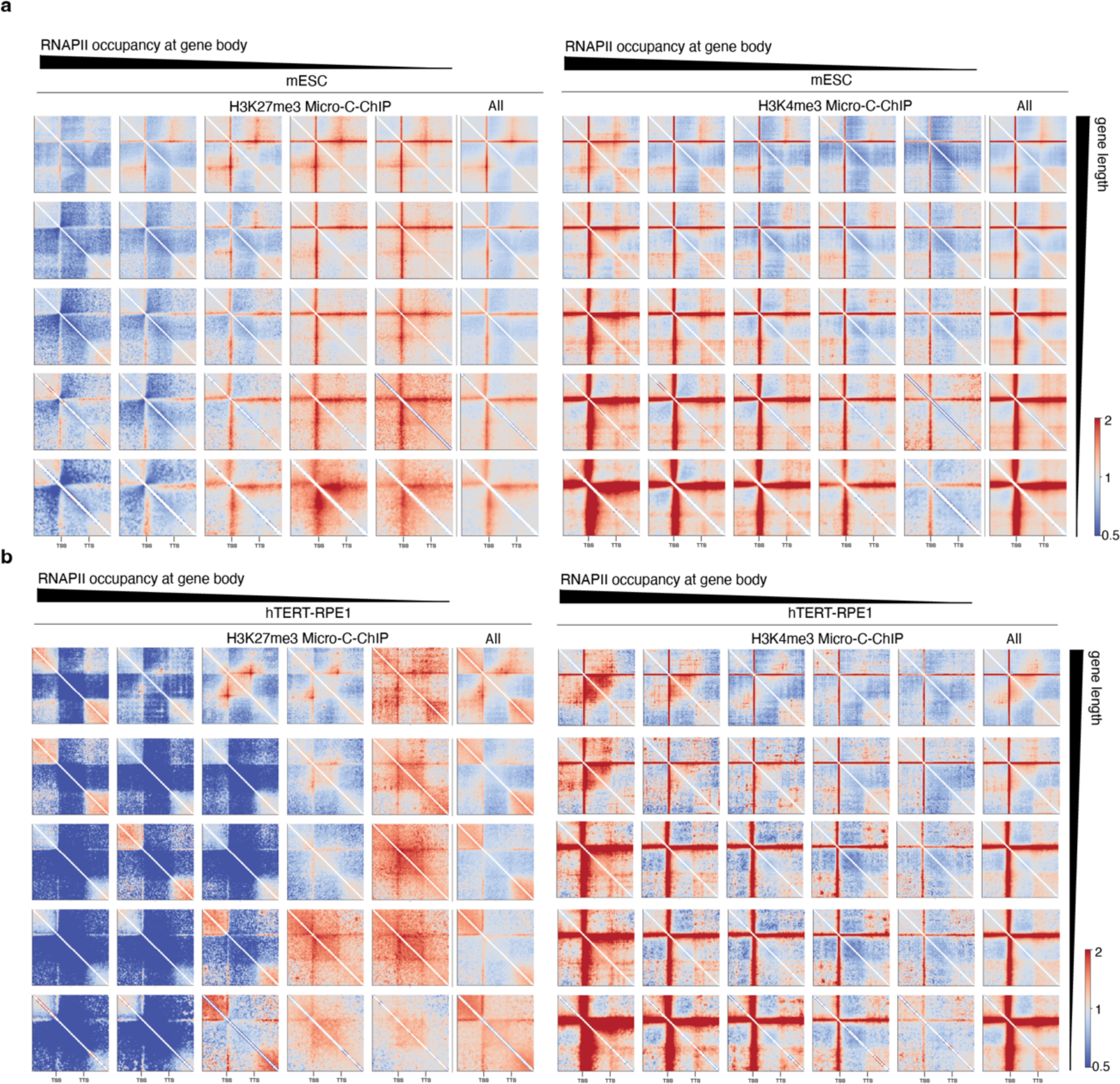
H3K27me3-associated stripes are hallmarks of transcriptionally inactive promoters in pluripotent cells. **a, b**, Pileup meta-gene plots around genes classified by length (vertical axis) and RNAPII occupancy (horizontal axis). The aggregated rescaled observed/expected contact maps are shown for H3K27me3-(left) and H3K4me3-(right) enriched datasets of mESC (a) and hTERT-RPE1 cells (b). Genes were first divided into quintiles based on their length, then RNAPII mean signal was computed at the gene bodies separately for each length group and, finally, gene length groups were divided into quantiles based on RNAPII enrichment. TSS and TTS locations are shown below the matrices.

Next, we analyzed the chromatin structure at all promoters, sorted by RNAPII signal, and plotted TSS centered on-diagonal pileups (**Fig. 4b**). In agreement with previous work^1^, the H3K4me3 Micro-C-ChIP signal strengthens with the increase of RNAPII occupancy in both cell types. However, the promoter-originating stripes are present even in the bottom-quantile RNAPII occupancy in mESC. In contrast, H3K27me3-enriched Micro-C-ChIP data shows a gradual decrease in interaction signal with increasing RNAPII occupancy in mESC, whereas, in hTERT-RPE1 cells, no H3K27me3 associated promoter-stripes are detectable in any cluster. Importantly, the finding that H3K27me3 data shows loop extrusion patterns in mESC but not in hTERT-RPE1 cells demonstrates that the detection of loop-extrusion patterns is not a methodological artifact of Micro-C-ChIP at H3K27me3 enriched promoters. Next, we corroborated these results by rescaling meta-gene pileup analysis stratifying genes by length and RNAPII occupancy (**Extended Data Fig. 4a, b**). We observed that the TSS- originating stripes associated with bottom RNAPII quantiles in the H3K27me3-enriched mESC dataset reach beyond the gene body independent of gene length (**Extended Data Fig. 4a**). In hTERT-RPE1 H3K27me3 associates with intra-gene compaction (**Extended Data Fig. 4b**). In both cell types, H3K4me3 Micro-C-ChIP shows TSS-associated stripes that scale with RNAPII occupancy, consistent with Fig. 4b, and reach beyond the transcription termination site. Our data suggests that H3K27me3 is part of a different chromatin architecture in mESC compared to hTERT-RPE1, and loop-extrusion features, such as stripes and chromosome loops, accompany this architecture.

### Bivalent promoter-connectome as a feature of chromatin organization in mESC

To get further insight into the commonalities of promoter-originating interactions in H3K4me3 and H3K27me3 Micro-C-ChIP datasets in mESC, we identified stripes with the *Stripenn* pipeline^37^. Pileup analysis at called 5’ and 3’ stripes in mESCs shows that H3K4me3 and H3K27me3 Micro-C-ChIP signals are enriched at stripes called on data from either mark (**Fig. 5a**), arguing that bivalent regions show loop-extrusion features in mESC. In contrast, H3K4me3-called stripes from hTERT-RPE1 cells did not show any enrichment in H3K27me3 Micro-C-ChIP data and the identification of stripes from H3K27me3 data with *Stripenn* did not yield reliable stripe calls (visual inspection, data not shown). To further dissect the bivalent nature of stripes common for both histone marks, we intersected stripe anchors of the same orientation (5’ in this instance) identified in H3K27me3 and H3K4me3 Micro-C-ChIP in mESC (**Fig. 5b**). Of 2170 H3K27me3 stripes, over 700 were shared with H3K4me3 stripes, while 1451 were unique for H3K27me3 Micro-C-ChIP. More than 70% of the shared stripes are TSS, the majority at which H3K27me3 and H3K4me3 peaks overlap. This confirms the presence of interactions originating from bivalent promoters.

**Fig. 5:**
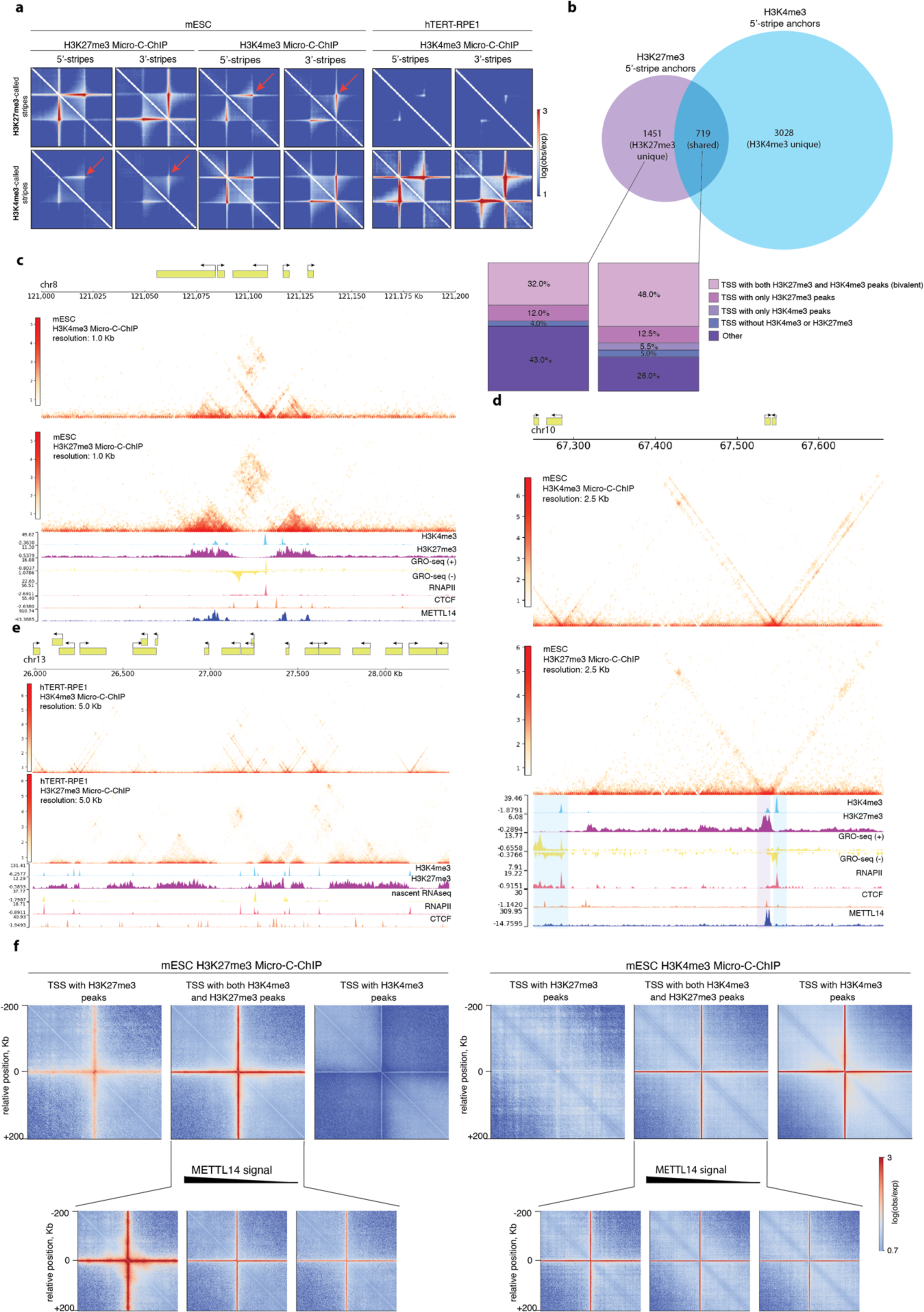
Chromatin organization in mESC is characterized by bivalent promoter-connectome. **a**, Rescaled pileup plots of 5’-and 3’-stripes at the mESC H3K27me3 (left) and H3K4me3 (middle) and hTERT-RPE1 H3K4me3 (right) Micro-C-ChIP averaged contact maps. The average intensity of interactions across stripes that were detected in the H3K27me3 (upper) and H3K4me3 (lower) Micro-C-ChIP datasets is shown at the corresponding and non-corresponding heatmaps. The red arrows point at the stripes that were called at the non-corresponding datasets in mESC. **b**, Venn diagram of unique and shared H3K4me3 or H3K27me3 Micro-C-ChIP unidirectional (5’ only) stripe anchors. H3K27me3-unique and shared stripe anchors were classified based on overlap with TSS locations (±500 bp). TSS have been categorized according to their 1D epigenetic status: with both H3K27me3 and H3K4me3 peaks (bivalent), with only H3K27me3 peaks, with only H3K4me3 peaks or without any histone marks. **c**, **d**, **e**, Interaction heatmaps of H3K4me3 (top) and H3K27me3 (bottom) in mESC (c, d) and hTERT-RPE1 (e) at 1-kb (c), 2.5-kb (d) and 5-kb (e) resolutions. Snapshots at c and d show common 3D organization pattern for datasets enriched for the different histone marks in the pluripotent cells, while interactions in hTERT-RPE1 are unique for both histone marks (e). 1D chromatin tracks are shown below the contact maps. **f**, On-diagonal pileup plots of TSS classified as in b. Aggregated contact maps are shown for H3K27me3 (left) and H3K4me3 (right) Micro-C-ChIP. TSS overlapping with both H3K27me3 and H3K4me3 ChIP-seq peaks have been further stratified by the METTL14 signal (bottom panel).

**Extended Data Fig. 5.**
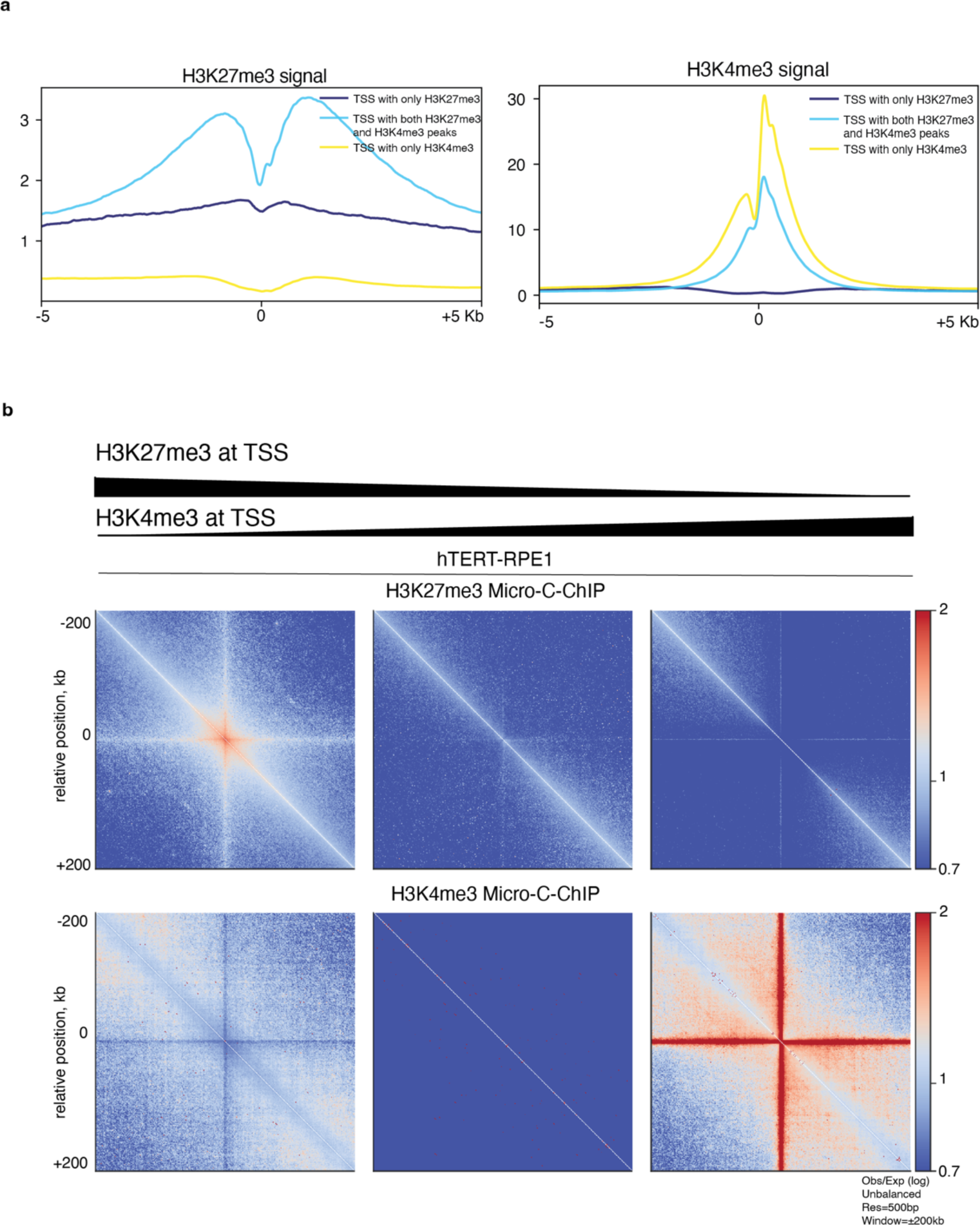
Bivalent promoter-associated interactions are not observed in hTERT-RPE1. **a**, Profile plots of H3K27me3 (left) and H3K4me3 (right) ChIP-seq signal at TSS classified as in Fig. 3b. The data is shown for ±5 kb window around TSS. **b**, On-diagonal pileup plots of TSS clustered based on H3K27me3 and H3K4me3 mean signal (left panel – high H3K27me3 and low H3K4me3, middle – medium H3K27me3 and low H3K4me3, right – low H3K27me3 and high H3K4me3). The aggregated contact maps around TSS are shown for H3K27me3 (top) and H3K4me3 (bottom) Micro-C-ChIP at ±200 kb window.

The architecture of bivalent chromatin remains a topic of debate^38^. Initially, bivalent promoters in *D. melanogaster* were found to be associated with paused RNAPII, leading to the idea that bivalent marks and paused RNAPII position promoters to respond to developmental queues readily^39,40^. However, careful GRO- and ChIP-seq experiments show that bivalent promoters at developmental genes are depleted of RNAPII^41^. Co-ChIP and Re-ChIP technologies allow to identify co-occurrence of H3K4me3 and H3K27me3^42–44^. Recent Re-ChIP experiments report around 8000 genes enriched for H3K4me3 and H3K27me3 at the same nucleosomes^44^. In addition, it was recently reported that bivalent regions in mESC are enriched in the RNA-binding protein METTL14, which regulates PRC2 and KDM5B localization to bivalent domains^45^. Indeed, visual inspection of mESC datasets confirms the enrichment of METTL14 at H3K4me3 and H3K27me3 shared regions (**Fig. 5c, d**). Furthermore, in contrast to pluripotent mouse cells, hTERT-RPE1 Micro-C-ChIP displays distinct patterns for histone marks and the absence of bivalent chromatin regions (**Fig. 5e**).

To investigate the architecture of bivalent promoters, we intersected TSS with H3K27me3 and H3K4me3 peaks and grouped them into three classes: H3K27me3 only, H3K27me and H3K4me3, and H3K4me3 only (**Fig. 5f, top panel**). We observed that the stripe signal for H3K27me3 Micro-C-ChIP is substantially stronger at promoters that are also marked by H3K27me3 and H3K4me3 compared to promoters that are only marked by H3K27me3 (**Fig. 5f**, **Extended Data Fig. 5a**). In contrast, the H3K4me3 Micro-C-ChIP signal is stronger for promoters that are only marked by H3K4me3 compared to promoters with both marks (**Fig. 5f**, right panel), presumably because these sites are associated with ongoing transcription.

Identifying bivalent promoters by two histone marks is complicated as it depends on data quality and signal distribution for peak calling. Thus, we sorted promoters with overlapping H3K4me3 and H3K27me3 peaks by the METTL14 signal to further select for bivalent promoters (**Fig. 5f**). It is noticeable that the interaction signal of H3K27me3 Micro-C-ChIP scales with METTL14 signal but the H3K4me3 signal remains similar. Visual inspection of mESC datasets complements the results of the 2D pileup analysis from Fig. 5f. It shows that bivalent TSS regions are characterized by common stripes for both H3K4me3 and H3K27me3 Micro-C-ChIP, overlapping ChIP-seq peak regions, enrichment of Metll14 and absence of RNAPII and ongoing transcription (**Fig. 5c, d**). Importantly, hTERT-RPE1 Micro-C-ChIP displays unique features for both histone marks and, therefore, no bivalency pattern is observed either via visual inspection or TSS-centered 2D pileup analysis (**Fig. 5e, Extended Data Fig. 5b**).

Micro-C-ChIP reveals that epigenetic states are reflected on chromatin folding patterns. And since the epigenetic landscape varies between cell types, differences result in distinct chromatin organization patterns. Moreover, enriching for H3K4me3 and H3K27me3 histone marks dissects the spatial organization of active and inactive chromatin. We show that there are not only contacts between transcriptionally active but also inactive promoters in pluripotent cells. Those promoters possess a bivalent nature where both H3K4me3 and H3K27me3 marks are present, resulting in the common promoter-associated stripes.

## Discussion

In this manuscript, we present a high-resolution epimark-specific enrichment strategy to measure 3-dimensional mammalian genome organization. Compared to bulk Micro-C, Micro-C-ChIP detects more short-range interactions indicating an improved signal-to-noise ratio. Furthermore, compared to other methods, such as MChIP-C that rely on formaldehyde-crosslinking only and that omits the enrichment of proximity-ligated fragments utilizing biotin-enrichment, our strategy yields a significantly higher fraction of informative 3D contacts^46^. At the level of individual genes, Micro-C-ChIP detects interactions that were, until now, only accessible through capture strategies, such as RCMC^23^ or TMCC^47^, or resource-intensive sequencing experiments. In contrast to RCMC and TMCC, Micro-C-ChIP is not limited to individual genomic regions but profiles 3D chromatin interactions across the entire genome. Overall, due to its inherent use of MNase as the enzyme to fragment genomes, Micro-C-ChIP is ideal for unraveling histone mark-specific 3D genome interactions.

Loop-extrusion stripes are a feature of active transcription^1,35,37^. Here, we show that transcriptionally inactive bivalently marked promoters adopt similar patterns. One-sided halt or pause of loop-extrusion leads to the emergence of stripes in interaction heatmaps^13,35^. It is demonstrated that the strength of stripe features correlates with transcription rates and RNAPII occupancy at promoters^1,48^. Here, we demonstrate that the H3K27me3-marked promoters that are depleted for RNAPII display stripe formation in mESC. This argues that loop-extrusion occurs at TSS independent of transcription. In differentiated hTERT-RPE1 cells, H3K27me3 stripes are not detectable, even at silenced H3K27me3-enriched promoters. This demonstrates that promoter-associated stripes are not an artifact of Micro-C-ChIP and that transcriptionally inactive promoter-associated stripes are a feature of mouse pluripotent stem cells. It is intriguing to speculate that bivalent marking preserves a more active-like promoter architecture while maintaining a repressed state.

Micro-C-ChIP reveals bivalent chromatin architecture in mESC. The function of bivalent promoter marking in ES cells remains a debate in the field^32^. Currently, the idea is that bivalent promoters are in a state that allows them to rapidly switch to an active or inactive state depending on environmental queues during differentiation^42,49,50^. Our data supports this idea. Previous work shows that bivalent promoters adopt an open chromatin configuration and are highly connected^36,49,51^. For more stringent identification of the bivalent promoters, we included METTL14 enrichment at those sites^45^. Here, we show that bivalent promoters display an overall similar architecture to active genes when enriched for H3K4me3 and H3K27me3, including loop-extrusion pattern, i.e., stripes and loops. These coincide with METTL14 binding and are independent of RNAPII binding. Furthermore, we identify similar patterns at bivalent promoters in H3K4me3 and H3K27me3 Micro-C-ChIP experiments. Micro-C measures the interactions between two nucleosomes in 3D space. In the case of TSS stripes at bivalent promoters, promoter-proximal nucleosomes, marked by either H3K4me3 or H3K27me3, interact with most nucleosomes within a given region across the cell population. Therefore, our data shows that H3K27me3 marked nucleosomes engage in open promoter architecture. This supports the model of truly bivalently marked nucleosomes carrying both marks, supported by re-ChIP experiments, instead of a composite of two distinct heterogeneous cell populations^42–44^.

Taken together, our study provides a novel experimental strategy to map histone mark-specific 3D genome organization with a high signal-to-noise ratio. Micro-C-ChIP allows the identification of extensive promoter-promoter interaction networks with moderate sequencing effort compared to bulk and other Micro-C-based immunoprecipitation strategies^1,21,46^. We demonstrated that Micro-C-ChIP allows the dissection of histone mark-specific promoter architecture, e.g., at bivalent genes. Micro-C-Chip can be readily adapted for other histone marks. Importantly, the moderate sequencing requirement of Micro-C-ChIP will open novel opportunities to investigate the dynamics of 3D genome organization, for example, in depletion studies of essential factors of the loop extrusion machinery.

## Acknowledgments

We thank Anja Groth, Joshua Brickman, and Olliver J. Rando for critical discussion. We thank Magali Michaut and Heike Wollmann from the CPR/reNEW Genomics Platform for sequencing support. M. M. is supported by the Novo Nordisk Foundation (NNF0069780). Research in the Krietenstein Laboratory is supported by the Lundbeck Foundation (R368-2021-1076). Research at CPR is supported by the Novo Nordisk Foundation (NNF14CC0001).

## Experimental procedure

### Cell culture and dual crosslinking

Mouse embryonic stem cells (mESC, E14JU cell line with a 129/Ola background, male)^52^ were cultured on gelatin-coated dishes (0.2%) in serum + LIF conditions at 37 °C with 5% CO2. The E14 medium (DMEM-GlutaMAX-pyruvate (Gibco #31966-021) supplemented with FBS (15%; Sigma-Aldrich #F0392), LIF (homemade), 1× nonessential amino acids (Gibco #11140-050), 1× penicillin/streptomycin (Gibco #15140-122) and 2-mercaptoethanol (0.1 µM; Gibco #31350010), sterile filtered) was changed daily and cells were passaged every second day at a 1:8-1:10 ratio.

Primary retinal epithelial hTERT-RPE1 cells (ATCC, #CRL-4000) were grown in DMEM-GlutaMAX-pyruvate containing 10% FBS and 1× penicillin/streptomycin at 37°C under 5% CO2 in 75cm^2^ flasks splitted at 70%–80% confluency.

To perform crosslinking, cells were detached using Trypsin-EDTA (Gibco #25200-056), washed once with 1x DPBS w/o Mg^2+^ and Ca^2+^ (ThermoFisher #14190144), and resuspended in 1x DPBS (1 mio cells/mL) for crosslinking with 1% Formaldehyde for 10 min with rotation at RT. The reaction was quenched by adding 2.5 M glycine to a final concentration of 0.25 M. For the second crosslinking, cells were washed with 1x DPBS and crosslinked with 3 mM EGS (ThermoFisher #21565) in 1x DPBS at 4 mio cells/mL for 40 min with rotation at RT. Crosslinking was quenched with glycine of 0.4 M final concentration for 10 min at RT. Cells were washed with 1x DPBS and flash-frozen in aliquots.

### Preparative MNase digestion

5-10 mio frozen cells were used for one preparative Micro-C-ChIP. Cells were resuspended in 1x DPBS with 1x BSA (NEB #B9000S) added prior to resuspension to reduce the stickiness of the cells to the tub walls. Upon 10 min incubation on ice, cells were centrifugated for 5 min at 10000x g, washed with 500 μl MB#1 buffer (10 mM Tris-HCl, pH 7.5, 50 mM NaCl, 5 mM MgCl_2_, 1 mM CaCl_2_, 0.2% IGEPAL CA-630 (Sigma-Aldrich #I8896), 1x protease inhibitor (Halt Protease Inhibitor Cocktail, EDTA-free (100X); ThermoFisher #78425)), collected by centrifugation (10000x g, 5 min). The derived nuclei were resuspended in 1000 μl MB#1 and splitted into five 200 μl aliquots which are equivalent of 1 mio cells. Chromatin was digested with MNase (5 U for 1 mio cells) for 10 min at 37°C. MNase (Wortington Biochem #LS004798) concentration was selected for each batch based on the prior titration to yield 70-90% mono-nucleosomes. The digestion was stopped by the addition of 0.5 M EGTA (Bioworld #405200081) to a 4 mM final concentration and incubation at 65°C for 10 min.

### DNA end processing and ligation

After MNase digestion, samples were pooled to a 5 mio cell-equivalent for further processing. If more than 5 mio cells are to be used, these samples can be processed in parallel. 5% of the sample was used as an MNase digestion input control.

The chromatin was centrifugated for 5 min at 10000x g, washed with 500 μl 1x T4 DNA Ligase Buffer (NEB #B0202A) and collected by centrifugation (10000x g, 5 min). The pellet was resuspended in 90 µL of freshly prepared Micro-C “Master Mix 1” (10 μl 10x T4 DNA Ligase Buffer, 75 μl ddH2O, 5 μl T4 PNK (NEB #M0201L)). After incubation for 15 min at 37 °C with shaking, 10 μl Large Klenow Fragment (NEB #M0210L) was added and the chromatin is incubated for 15 min at 37°C with shaking. 100 µL of freshly prepared Micro-C “Master Mix 2” (10 μl 1 mM Biotin-dATP (Jenna Bioscience #NU-835-Bio14-S), 10 μl 1 mM Biotin-dCTP (Jenna Bioscience #NU-809-BioX-S), 1 μl 10 mM mix of dGTP and dTTP (NEB #N0442S, #N0443S), 5 μl 10x T4 DNA Ligase Buffer, 0.25 μl BSA, 23.75 μl ddH2O) was added. After incubation for 45 min at 25°C with shaking, the enzymatic reaction was quenched by adding 0.5 M EDTA to a final concentration of 30 mM and incubation for 20 min at 65°C with shaking. The chromatin was centrifugated (10000x g, 5 min), washed in 500 μl 1x T4 DNA Ligase Buffer, and collected by centrifugation (10000x g, 5 min). The chromatin pellet was resuspended in 500 μL of ligation reaction buffer (50 μl 10x T4 DNA Ligase Buffer, 2.5 µL BSA, 25 µL 400 U/µL T4 DNA Ligase, 422.5 µL ddH2O) and incubated for 2.5 hours at RT with slow rotation. After proximity ligation, the chromatin was collected by centrifugation (12000x g, 5 min), resuspended in 200 μl 1x NEBuffer 1 (NEB #B7001S) and 200 U NEB Exonuclease III (NEB #M0206S), and incubated for 15 min at 37°C to remove biotin from unligated ends.

### Solubilization and immunoprecipitation

After ExoIII digestion, chromatin was resuspended in the sonication buffer (0.1% IGEPAL CA-630, 50 mM NaCl, 10 mM Tris–HCl, pH 7.5, 5mM MgCl_2_, 2mM EDTA, 0.5% SDS (Invitrogen #15553-035)), 1x protease inhibitor) and sonicated for 10 cycles (sonication cycle: 30 sec ON, 30 sec OFF) using Bioruptor Plus (Diagenode). The samples were centrifugated at 14000x g for 10 minutes at 4°C and the supernatant containing the soluble chromatin was transferred to a fresh low-binding tube. At this stage, it is possible to pool 5 mio cell-equivalent aliquots, processed in parallel, to increase the immunoprecipitation yield. To control the efficient solubilization and to quantify DNA yield, 5-10% supernatant input was taken and visualized by 1.5% agarose gel electrophoresis. The supernatant was diluted 1:4 with ChIP dilution buffer (1.1% Triton X-100, 16.7 mM Tris-HCl, pH 7.5, 1.2 mM EDTA, 167 mM NaCl, 1x protease inhibitor). Anti-H3K27me3 (Cell Signaling Technology #9733s) and anti-H3K4me3 (Cell Signaling Technology #9751S) antibodies were added at concentration 10 μl of antibody per 10 μg of chromatin (typically 10-20 μl of antibody for 5-10 mio cells). Immunoprecipitation was performed overnight at 4° C with rotation. Magnetic beads (Pierce Protein A Magnetic Beads; ThermoFisher #88846) were added and incubated for 2 h at 4° C. The beads were washed in the following order: twice with low-salt wash buffer (20 mM Tris-HCl, pH 8.0, 150 mM NaCl, 2 mM EDTA, 1% Triton X-100, 0.1% SDS, 1x protease inhibitor), twice with high-salt wash buffer (20 mM Tris-HCl, pH 8.0, 500 mM NaCl, 2 mM EDTA, 1% Triton X-100, 0.1% SDS, 1x protease inhibitor), once with LiCl wash buffer (10 mM Tris–Cl, pH 8.0, 250 mM LiCl, 1% IGEPAL, 1% Sodium deoxycholate, 1mM EDTA, 1x protease inhibitor) and twice with 1X TE buffer (10 mM Tris-HCl, pH 8, 1 mM EDTA). After the last wash, antibody-protein-DNA complexes were eluted from the beads and reverse-crosslinked by adding 250 μl Elution buffer (20 mM Tris-HCl, pH 8.0, 10 mM EDTA, 5 mM EGTA, 300 mM NaCl, 1% SDS, Proteinase K, ddH2O) and incubated for at least 2 h at 65°C. DNA was purified via Phenol:Chloroform:Isoamylalcohol (Invitrogen #15593-031) extraction followed by ethanol precipitation and resuspended in 50 µL 0.1x TE buffer (1 mM Tris-HCl, pH 7.5, 0.1 mM EDTA).

### Streptavidin pull-down and on-bead library preparation

10 μl Dynabeads™ MyOne™ Streptavidin C1 beads (Invitrogen #65001) were washed twice with 500 μl 1x TBW (5 mM Tris-HCl, pH 7.5, 0.5 mM EDTA, 1 M NaCl) and resuspended in 150 μl 2x TBW (10 mM Tris-HCl, pH 7.5, 1 mM EDTA, 2 M NaCl). The resuspended beads and 100 µL 0.1x TE were added to the 50 µL sample and incubated with rotation at RT for 20 min. The beads were washed twice with 500 μl 1x TBW, once with 300 μl 0.1x TE buffer, resuspended in 50 μl 0.1x TE buffer and transferred to the PCR tubes. Sequencing libraries were prepared with NEBNext® Ultra II DNA Library Prep Kit for Illumina® (NEB #E7645). Note: because the sample remains attached to beads, all purification and size selection steps were omitted. Instead, the beads were washed twice with 300 μl 1x TBW and once with 300 μl 0.1x TE and resuspended in 20 μl 0.1x TE. PCR master mix (64 µL of H2O, 100 µL of Q5 high-fidelity DNA polymerase (NEB #M0544S), 8 µL of i5 primer, 8 µL of i7 primer (NEBNext Multiplex Oligos for Illumina (Dual Index primers); NEB #E7600S) was added to the sample. The reaction mixture was split into 100 µL aliquots. PCR amplification was performed for 14-16 cycles according to NEBNext Ultra II DNA library prep kit for Illumina. After the PCR reaction, DNA was purified with SPRI size-selection beads (Beckman Coulter #B23319) at a ratio of 1:0.9 according to the manufacturer’s protocol and eluted in 20 µL of 0.1x TE.

### Sequencing

The samples were sequenced on an Illumina HiSeq 2000. We used Illumina 50 bp paired-end sequencing to obtain ∼200-300 million total reads for each replicate in this study (**Extended Data Fig. 1b**).

## Data analysis

### Mapping

Micro-C-ChIP paired-end reads were processed with the distiller pipeline (https://github.com/mirnylab/distiller-nf). Briefly, reads were mapped to the reference genome (mm10 and hg38 as mouse and human reference assemblies, respectively) via bwa mem with flags –SP (v0.7.17). Alignments were parsed and .pairs files were generated using the pairtools package^53^ (v0.3.0). PCR/optical duplicates were filtered with pairtools dedup function with “-max-mismatch 1”. Valid pairs with high mapping quality scores on both sides (MAPQ > 30) were aggregated into binned matrices of Micro-C-ChIP interactions using the cooler^54^ (v0.9.2) package into multiresolution cooler (mcool) files (100 bp, 200 bp, 500 bp, 1 kb, 2.5 kb, 5 kb, 10 kb, 25 kb, 50 kb, 100 kb, 500 kb, 1 Mb). All contact matrices displayed throughout the manuscript are unbalanced (**Fig. 1e, Fig. 2e, Fig. 3a, Fig. 5c, d, e, Extended Data Fig. 1c, d, 2a, b**). The reproducibility of Micro-C-ChIP replicates was evaluated by HiCRep^55^ (v0.2.6) for contact maps at 10 kb resolution (**Extended Data Fig. 1a**).

### Visualization

Contact maps shown in figures alongside gene annotations and genomic tracks (ChIP-seq, GRO-seq, nascent RNA-seq) were generated with CoolBox^56^ (v0.3.9). The multi-cooler files used for contact map visualization were generated with cooler cload pairs and cooler zoomify from the valid pairs files. The valid pairs reads were shifted by 73 bp with respect to the read orientation. This was done to account for the approximate locations of the nucleosome dyad axes. Additionally, the valid pairs separated by less than 500 bp were disregarded to remove reads from undigested di-nucleosomal contaminants.

### Analysis of publicly available ChIP-seq datasets

All the ChIP-seq datasets that were downloaded as raw fastq files (see Supplementary Table 1) were reanalyzed. Bowtie2 (v2.4.2) was used for mapping the fastq files to the hg38 or mm10 reference genome. The signal tracks were produced via deeptools bamCoverage (v3.5.1) with the parameters “-binSize 20 -normalizeUsing RPKM - smoothLength 60 -extendReads 150 -centerReads”.

### 1D analysis

We used the TSS (transcription start sites) locations of the mm10 and hg38 refTSS annotations^57^ with a ±500 bp window for defining promoter regions. Enhancers were defined based on the overlap of publicly available ChIP-seq datasets for H3K4me1 (ENCFF440FYE for mESC, GSE176035 for hTERT-RPE1) and H3K27ac (ENCFF194TQD for mESC, GSE113399 for hTERT-RPE1) excluding regions that overlap with promoters. Putative enhancer-promoter (E-P) pairs separated by <100 kb distance were created with a custom R script and putative promoter-promoter (P-P) pairs (80-100 kb distance) were generated with the bioframe^58^ (v0.4.1) pair-by-distance function (**Fig. 1g, 2c)**.

CTCF motifs were obtained from JASPAR (JASPAR motif ID #MA0139.1) and overlapped with the CTCF ChIP-seq peaks (GSE90994 for mESC, GSE176035 for hTERT-RPE1) using bioframe^58^ (v0.4.1). CTCF ChIP-seq peaks that are associated with the strongest motif underlying that peak were then overlapped with the SMC1A peaks (GSE123636 for mESC, GSE146766 for hTERT-RPE1) to obtain cohesin-bound CTCF sites.

We stratified TSS by RNAPII occupancy calculating the mean RNAPII ChIP-seq signal (GSE52071 for mESC, GSE201982 for hTERT-RPE1) in a ±500-bp window around all TSS locations and divided them into quantiles (**Fig. 4c, d**). TSS locations were also stratified based on the overlap with H3K27me3 and/or H3K4me3 peaks (Supplementary Table 1) using bedtools^59^ (v2.30.0) intersect (**Fig. 5b, f**). To stringently identify bivalent TSS, we further stratified the TSS where H3K27me3 and H3K4me3 peaks overlap by METTl14 occupancy and divided into three quantiles where the top METTl14 quantile corresponds to bivalent promoters (**Fig. 5f**).

Average signal plots for Micro-C-ChIP datasets were generated with deeptools^60^ (v3.5.1). The ChIP-seq peaks for a corresponding histone mark were used as alignment points. The signal around the alignment points was averaged within a ±30 kb window. The Micro-C-ChIP signal tracks used for the signal tracks and average signal visualization were generated with bamCoverage using CPM normalization and 20 bp bin size (**Fig. 1d, 2c, Extended Data Fig. 1c**). For H3K27me3 and H3K4me3 ChIP-seq signal (GSE154379 and ENCFF806NDV, respectively) in mESC, metaplots over TSS, which were stratified based on the overlap with H3K27me3 and/or H3K4me3 peaks, were generated using a ±5 kb window (**Extended Data Fig. 5a**).

### Loop detection and anchor analysis

The loops in H3K4me3 Micro-C-ChIP datasets from mESC and hTERT-RPE1 cells were predicted with Peakachu^61^ (v2.2) using a pretrained model (RNAPII ChIA-PET dataset in WTC11 cells) offered in the pipeline. The loops were called at the 2.5- and 5-kb resolution unbalanced contact maps (**Fig. 3a**).

The loops were visualized using Coolbox. To classify loops, we intersected loop anchors with the promoter, enhancer, or cohesin-bound CTCF regions (**Fig. 3b**). CTCF-classified anchors do not include regions that overlap promoter or enhancer regions. Anchors which do not overlap any of three features were defined as “Other”. The promoter-overlapping anchors were further classified based on the overlap with the cohesin-bound CTCF regions.

Anchor valency was used as a proxy of loop engagement of one anchor (**Fig. 3e**). For this, the genomic coordinates of the left and the right anchors for each loop were concatenated resulting in 1D anchor list. We then used bedtools merge with the parameters “-c 1 -o count” computing the number of merged anchors in a given genomic locus.

The number of interactions per anchor and the lengths of the loops were visualized in R (v 4.1.2) using ggplot2 (v3.4.2) (**Fig. 3c, e**).

### Stripe calling

The stripes were identified with Stripenn^37^ (v1.1.65.18) at 2.5- and 5-kb resolution contact maps (**Fig. 5a**). The called stripes were classified as 5’- and 3’-stripes having anchors at 5′- or 3’-end of the stripe domains, respectively.

### Pileup analysis

Contact maps used for the pileup plots were unbalanced throughout the manuscript. To account for distance decay in the pileups, all pileups demonstrate aggregated observed-over-expected contact maps where expected interactions for chromosome arms were calculated beforehand. The colorbars to the side from the pileups show the averaged observed-over-expected signal in logarithmic scale.

The off-diagonal pileup analysis was done to calculate Micro-C-ChIP contact aggregation at paired ChIP-seq peaks obtained from the publicly available datasets (**Fig. 1f, 2b**) using cooltools^62^ (v0.5.1). First, the strongest ChIP-seq peaks were intersected using bioframe^58^(v0.4.1) to create paired sites separated by the distance between 10 and 300 kb. The paired ChIP-seq peaks were piled up on the center of 500 bp resolution maps around ±30 kb window. 500-bp resolution was used for the pileup plots.

The off-diagonal pileup analysis of P-P loops that were identified in H3K4me3 Micro-C-ChIP and further stratified by RNAPII signal (**Fig. 3d, 4a, b**) was performed with Coolpup^63^ (v1.1.0) leveraging the advantage of subsetting the snippets. 100-kb flank was used for the pileup analysis of P-P loops for 500-bp resolution Micro-C-ChIP matrices (**Fig. 3d**) and 20-kb flank for the pileup analysis of P-P loops with anchors stratified by RNAPII level for 100-bp resolution matrices (**Fig. 4a, b**).

The putative E-P/P-P pairs were centered and piled up at 200-bp resolution matrices around ±10 kb window (**Fig. 1g, 2d**).

Instead of aggregating at the intersection of ChIP-seq peaks or loop anchors, we applied the same approach to analyze the genome-wide target-centered contact intensity (**Fig. 4c, d, 5a, f, Extended Data Fig. 4a, b, 5b**). All on-diagonal (local) pileup maps were computed with Coolpup (v1.1.0). TSS-centered pileups were plotted around ±200 kb window where the TSS coordinates were stratified based on the RNAPII mean signal (**Fig. 4c, d**), the overlap with the H3K27me3 and/or H3K4me3 ChIP-seq peaks and METTL14 mean signal (**Fig. 5f**).

The rescaled local pileup analysis was applied to evaluate average interaction signal across the identified 5’-and 3’-stripes (**Fig. 5a**) and gene body (**Extended Data Fig. 4a, b**) using “-rescale” parameter with rescale flank equal 1 rescaling the pileups to the same shape and size. With regards to the pileup analysis of gene folding, genes were first grouped based on their length, then mean RNAPII signal was computed at the gene bodies (from TSS to TTS (transcription termination sites)) separately for each length group. Finally, each length group was divided into quantiles based on RNAPII enrichment and rescaled to the same length from TSS to TTS on the x axis. TSS and TTS locations are shown below the matrices. mm10 and hg38 RefSeq-derived gene coordinates^64^ were used in this analysis.

## References

1. Hsieh, T.-H. S. et al. Resolving the 3D Landscape of Transcription-Linked Mammalian Chromatin Folding. Mol. Cell 78, 539–553.e8 (2020).

2. Hsieh, T.-H. S. et al. Enhancer–promoter interactions and transcription are largely maintained upon acute loss of CTCF, cohesin, WAPL or YY1. Nat. Genet. 54, 1919–1932 (2022).

3. Barshad, G. et al. RNA polymerase II dynamics shape enhancer–promoter interactions. Nat. Genet. 55, 1370–1380 (2023).

4. Zhang, S., Übelmesser, N., Barbieri, M. & Papantonis, A. Enhancer–promoter contact formation requires RNAPII and antagonizes loop extrusion. Nat. Genet. 55, 832–840 (2023).

5. Banigan, E. J. et al. Transcription shapes 3D chromatin organization by interacting with loop extrusion. Proc. Natl. Acad. Sci. 120, e2210480120 (2023).

6. Sanders, J. T. et al. Radiation-induced DNA damage and repair effects on 3D genome organization. Nat. Commun. 11, 6178 (2020).

7. Arnould, C. et al. Chromatin compartmentalization regulates the response to DNA damage. Nature 1–10 (2023) doi:10.1038/s41586-023-06635-y.

8. Marchal, C., Sima, J. & Gilbert, D. M. Control of DNA replication timing in the 3D genome. Nat. Rev. Mol. Cell Biol. 20, 721–737 (2019).

9. Lieberman-Aiden, E. et al. Comprehensive Mapping of Long-Range Interactions Reveals Folding Principles of the Human Genome. Science 326, 289–293 (2009).

10. Nora, E. P. et al. Spatial partitioning of the regulatory landscape of the X-inactivation centre. Nature 485, 381–385 (2012).

11. Dixon, J. R. et al. Topological domains in mammalian genomes identified by analysis of chromatin interactions. Nature 485, 376–380 (2012).

12. Fudenberg, G. et al. Formation of Chromosomal Domains by Loop Extrusion. Cell Rep. 15, 2038–2049 (2016).

13. Fudenberg, G., Abdennur, N., Imakaev, M., Goloborodko, A. & Mirny, L. A. Emerging Evidence of Chromosome Folding by Loop Extrusion. Cold Spring Harb. Symp. Quant. Biol. 82, 45–55 (2017).

14. Nora, E. P. et al. Targeted Degradation of CTCF Decouples Local Insulation of Chromosome Domains from Genomic Compartmentalization. Cell 169, 930–944.e22 (2017).

15. Rao, S. S. P. et al. A 3D Map of the Human Genome at Kilobase Resolution Reveals Principles of Chromatin Looping. Cell 162, 687–688 (2015).

16. Rao, S. S. P. et al. Cohesin Loss Eliminates All Loop Domains. Cell 171, 305–320.e24 (2017).

17. Taatjes, D. J., Marr, M. T. & Tjian, R. Regulatory diversity among metazoan co-activator complexes. Nat. Rev. Mol. Cell Biol. 5, 403–410 (2004).

18. Oksuz, B. A. et al. Systematic evaluation of chromosome conformation capture assays. Nat. Methods 18, 1046–1055 (2021).

19. Hsieh, T.-H. S. et al. Mapping Nucleosome Resolution Chromosome Folding in Yeast by Micro-C. Cell 162, 108–119 (2015).

20. Hsieh, T.-H. S., Fudenberg, G., Goloborodko, A. & Rando, O. J. Micro-C XL: assaying chromosome conformation from the nucleosome to the entire genome. Nat. methods 13, 1009–1011 (2016).

21. Krietenstein, N. et al. Ultrastructural Details of Mammalian Chromosome Architecture. Mol. Cell 78, 554–565.e7 (2020).

22. Hsieh, T.-H. S. et al. Resolving the 3D Landscape of Transcription-Linked Mammalian Chromatin Folding. Mol Cell (2020) doi:10.1016/j.molcel.2020.03.002.

23. Goel, V. Y., Huseyin, M. K. & Hansen, A. S. Region Capture Micro-C reveals coalescence of enhancers and promoters into nested microcompartments. Nat. Genet. 55, 1048–1056 (2023).

24. Mumbach, M. R. et al. HiChIP: efficient and sensitive analysis of protein-directed genome architecture. Nat. Methods 13, 919–922 (2016).

25. Li, G. et al. ChIA-PET tool for comprehensive chromatin interaction analysis with paired-end tag sequencing. Genome Biol. 11, R22 (2010).

26. Fullwood, M. J. et al. An oestrogen-receptor-alpha-bound human chromatin interactome. Nature 462, 58–64 (2009).

27. Santos-Rosa, H. et al. Active genes are tri-methylated at K4 of histone H3. Nature 419, 407–411 (2002).

28. Guenther, M. G., Levine, S. S., Boyer, L. A., Jaenisch, R. & Young, R. A. A Chromatin Landmark and Transcription Initiation at Most Promoters in Human Cells. Cell 130, 77–88 (2007).

29. Wang, H. et al. H3K4me3 regulates RNA polymerase II promoter-proximal pause-release. Nature 615, 339–348 (2023).

30. Boyer, L. A. et al. Polycomb complexes repress developmental regulators in murine embryonic stem cells. Nature 441, 349–353 (2006).

31. Juan, A. H. et al. Roles of H3K27me2 and H3K27me3 Examined during Fate Specification of Embryonic Stem Cells. Cell Rep. 17, 1369–1382 (2016).

32. Bernstein, B. E. et al. A Bivalent Chromatin Structure Marks Key Developmental Genes in Embryonic Stem Cells. Cell 125, 315–326 (2006).

33. Azuara, V. et al. Chromatin signatures of pluripotent cell lines. Nat. Cell Biol. 8, 532–538 (2006).

34. Hawkins, R. D. et al. Distinct Epigenomic Landscapes of Pluripotent and Lineage-Committed Human Cells. Cell Stem Cell 6, 479–491 (2010).

35. Vian, L. et al. The Energetics and Physiological Impact of Cohesin Extrusion. Cell 173, 1165–1178.e20 (2018).

36. Schoenfelder, S. et al. Polycomb repressive complex PRC1 spatially constrains the mouse embryonic stem cell genome. Nat. Genet. 47, 1179–1186 (2014).

37. Yoon, S., Chandra, A. & Vahedi, G. Stripenn detects architectural stripes from chromatin conformation data using computer vision. Nat. Commun. 13, 1602 (2022).

38. Voigt, P., Tee, W.-W. & Reinberg, D. A double take on bivalent promoters. Genes Dev. 27, 1318–1338 (2013).

39. Muse, G. W. et al. RNA polymerase is poised for activation across the genome. Nat. Genet. 39, 1507–1511 (2007).

40. Zeitlinger, J. et al. RNA polymerase stalling at developmental control genes in the Drosophila melanogaster embryo. Nat. Genet. 39, 1512–1516 (2007).

41. Williams, L. H. et al. Pausing of RNA Polymerase II Regulates Mammalian Developmental Potential through Control of Signaling Networks. Mol. Cell 58, 311– 322 (2015).

42. Mas, G. et al. Promoter bivalency favors an open chromatin architecture in embryonic stem cells. Nat. Genet. 50, 1452–1462 (2018).

43. Weiner, A. et al. Co-ChIP enables genome-wide mapping of histone mark co-occurrence at single-molecule resolution. Nat. Biotechnol. 34, 953–961 (2016).

44. Ho, W., Seneviratne, J. A., Glancy, E. & Eckersley-Maslin, M. A. A low-input high resolution sequential chromatin immunoprecipitation method captures genome-wide dynamics of bivalent chromatin. bioRxiv 2023.09.18.558170 (2023) doi:10.1101/2023.09.18.558170.

45. Mu, M. et al. METTL14 regulates chromatin bivalent domains in mouse embryonic stem cells. Cell Rep. 42, 112650 (2023).

46. Golov, A. K., Gavrilov, A. A., Kaplan, N. & Razin, S. V. A genome-wide nucleosome-resolution map of promoter-centered interactions in human cells corroborates the enhancer-promoter looping model. bioRxiv 2023.02.12.528105 (2023) doi:10.1101/2023.02.12.528105.

47. Oudelaar, A. M. et al. Dynamics of the 4D genome during in vivo lineage specification and differentiation. Nat. Commun. 11, 2722 (2020).

48. Salari, H., Fourel, G. & Jost, D. Transcription regulates the spatio-temporal dynamics of genes through micro-compartmentalization. bioRxiv 2023.07.18.549489 (2023) doi:10.1101/2023.07.18.549489.

49. Jadhav, U. et al. Acquired Tissue-Specific Promoter Bivalency Is a Basis for PRC2 Necessity in Adult Cells. Cell 165, 1389–1400 (2016).

50. Zhang, J. et al. Highly enriched BEND3 prevents the premature activation of bivalent genes during differentiation. *Sci. (N. York*, NY) 375, 1053–1058 (2022).

51. Schoenfelder, S. et al. The pluripotent regulatory circuitry connecting promoters to their long-range interacting elements. Genome Res. 25, 582–597 (2015).

52. Hamilton, W. B. & Brickman, J. M. Erk Signaling Suppresses Embryonic Stem Cell Self-Renewal to Specify Endoderm. Cell Rep. 9, 2056–2070 (2014).

53. Open2C et al. Pairtools: from sequencing data to chromosome contacts. bioRxiv : Prepr. Serv. Biol. 2023.02.13.528389 (2023) doi:10.1101/2023.02.13.528389.

54. Abdennur, N. & Mirny, L. A. Cooler: scalable storage for Hi-C data and other genomically labeled arrays. Bioinformatics 36, 311–316 (2019).

55. Yang, T. et al. HiCRep: assessing the reproducibility of Hi-C data using a stratum-adjusted correlation coefficient. Genome Res. 27, 1939–1949 (2017).

56. Xu, W. et al. CoolBox: a flexible toolkit for visual analysis of genomics data. BMC Bioinform. 22, 489 (2021).

57. Abugessaisa, I. et al. refTSS: A Reference Data Set for Human and Mouse Transcription Start Sites. J. Mol. Biol. 431, 2407–2422 (2019).

58. Open2C et al. Bioframe: Operations on Genomic Intervals in Pandas Dataframes. bioRxiv 2022.02.16.480748 (2022) doi:10.1101/2022.02.16.480748.

59. Quinlan, A. R. & Hall, I. M. BEDTools: a flexible suite of utilities for comparing genomic features. Bioinformatics 26, 841–842 (2010).

60. Ramírez, F. et al. deepTools2: a next generation web server for deep-sequencing data analysis. Nucleic Acids Res. 44, W160–W165 (2016).

61. Salameh, T. J. et al. A supervised learning framework for chromatin loop detection in genome-wide contact maps. Nat. Commun. 11, 3428 (2020).

62. Open2C et al. Cooltools: enabling high-resolution Hi-C analysis in Python. bioRxiv 2022.10.31.514564 (2022) doi:10.1101/2022.10.31.514564.

63. Flyamer, I. M., Illingworth, R. S. & Bickmore, W. A. Coolpup.py: versatile pile-up analysis of Hi-C data. Bioinformatics 36, 2980–2985 (2020).

64. O’Leary, N. A. et al. Reference sequence (RefSeq) database at NCBI: current status, taxonomic expansion, and functional annotation. Nucleic Acids Res. 44, D733–D745 (2016).

